# DeepMHCII: A Novel Binding Core-Aware Deep Interaction Model for Accurate MHC II-peptide Binding Affinity Prediction

**DOI:** 10.1101/2021.12.27.474242

**Authors:** Ronghui You, Wei Qu, Hiroshi Mamitsuka, Shanfeng Zhu

## Abstract

Computationally predicting MHC-peptide binding affinity is an important problem in immunological bioinformatics. Recent cutting-edge deep learning-based methods for this problem are unable to achieve satisfactory performance for MHC class II molecules. This is because such methods generate the input by simply concatenating the two given sequences: (the estimated binding core of) a peptide and (the pseudo sequence of) an MHC class II molecule, ignoring the biological knowledge behind the interactions of the two molecules. We thus propose a binding core-aware deep learning-based model, DeepMHCII, with binding interaction convolution layer (BICL), which allows integrating all potential binding cores (in a given peptide) and the MHC pseudo (binding) sequence, through modeling the interaction with multiple convolutional kernels. Extensive empirical experiments with four large-scale datasets demonstrate that DeepMHCII significantly outperformed four state-of-the-art methods under numerous settings, such as five-fold cross-validation, leave one molecule out, validation with independent testing sets, and binding core prediction. All these results with visualization of the predicted binding cores indicate the effectiveness and importance of properly modeling biological facts in deep learning for high performance and knowledge discovery. DeepMHCII is publicly available at https://weilab.sjtu.edu.cn/DeepMHCII/.

## 1 Introduction

Major Histocompatibility Complex (MHC) molecules play a significant role in the T-cell mediated adaptive immune response [1]. MHC molecules first bind peptide fragments derived from pathogens, and then present the peptides to the surface of antigen presenting cells (APC). After the MHC-peptide complexes are recognized by T-cell receptors (TCR), adaptive immune response will be stimulated to fight against and eliminate invading pathogens. Accurate identification of MHC binding peptides is thus crucial for not only elucidating the mechanism of immune recognition, but also facilitating the design of of peptide-based vaccine and cancer immunotherapy [2]. As biochemical experiments are time consuming and labor intensive, computational approaches for predicting MHC binding peptides have become increasingly important and have been utilized to prioritize a small number of promising candidates for further verification by biochemical experiments [3].

There are two major classes of MHC molecules: MHC class I (MHC-I) and MHC class II (MHC-II). MHC-I molecules have one chain (*α*) and MHC-II molecules have two chains (*α* and *β*). Human MHC-II molecules are encoded in the human leucocyte antigen (HLA) gene complex involving three types of molecules: DP, DQ, DR, while mouse MHC-II are encoded in the histocompatibility 2 (H-2) [4]. MHC-I and MHC-II molecules play different roles in adaptive immune response. MHC-I molecules bind a rather fixed length of short peptides (usually 8-11 amino acids) from endogenous antigen, and these peptides are presented to cytotoxic T lymphocytes (CTL). In contrast, MHC-II molecules bind a more diverse length of peptides from exogenous antigen, and these peptides are presented to helper T lymphocytes. It has also been reported that MHC-II peptide binding involves the B-cell mediated adaptive immune response. From these aspects, predicting peptides binding to MHC-II is more challenging than that of MHC-I [5]. Currently the state-of-the-art methods for MHC-I peptide binding prediction can achieve an Area Under the ROC Curve (AUC) of around 0.9, while AUC by the prediction methods for MHC-II is usually unable to reach 0.8, particularly for MHC-II molecules with very few or no binding peptide data [6, 7], which is far from satisfactory. Moreover, quantitative prediction of MHC peptide binding is more useful in practice than binary classification for selecting a small number of promising candidate peptides. It is thus imperative to develop accurate computational methods for MHC-II peptide binding affinity prediction.

The challenges come from two sides: peptides and MHC-II molecules. For the peptide side, the binding groove of MHC-II molecules is open at both ends, which causes large variation on the length of binding peptides, ranging from 10 to 30 amino acids (typically 12-16). The binding groove of MHC-II has nine pockets, where one amino acid residue of the binding core of a binding peptide fits to one pocket normally [1]. The peptide-MHC binding affinity is primarily determined by the interaction between MHC-II molecules and the binding core of peptides. However it has been found that peptides flanking regions (PFRs) which is outside of the binding core also affect the binding affinity [8, 9]. Thus there are two issues for the peptide side: 1) the flexibility in length and 2) the location of the binding core. For the MHC-II molecule side, MHC-II molecules are highly polymorphic. There are thousands of MHC-II molecules and each MHC-II molecule has its own binding specificity. Also MHC molecules are proteins to be represented by amino acid sequences with longer and more diverse lengths than peptides. In addition currently only dozens of MHC-II molecules have hundreds of binding affinity data in Immune epitope database and analysis resource (IEDB) [10], and a vast majority of MHC-II molecules have very few or even no binding data. Thus there are three issues for the MHC-II side: 1) thousands of MHC-II molecule with different binding specificity, 2) long and size-flexible sequences and 3) data scarcity for most of MHC-II molecules. These aspects have made it difficult to model the interaction of MHC-II peptide binding accurately.

Computational methods for MHC-II peptide binding prediction can be divided into two categories: allele-specific and pan-specific [7]. Allele-specific methods can predict only MHC-II molecules in the training set, while pan-specific methods can predict MHC-II molecules even with no training data, which is thus the focus of our work. Traditionally, pan-specific methods have been developed by various different techniques, such as position specific scoring matrices (PSSM) [11], artificial neural network (ANN) [6] and kernel based methods [12]. The most established method is NetMHCIIpan (latest version for MHC-II peptide binding prediction is NetMHCIIpan-3.2 [6]), an ANN-based method, which pioneered using pseudo sequence to represent an MHC-II molecule. ANN deals with only a fixed sized input, and so NetMHCIIpan first estimates the binding core in a given peptide and then trains an ANN using the estimated binding cores and the pseudo sequences. The whole process is repeated until convergence. However, the first estimation of the binding core might be inaccurate, which affects the predictive performance heavily. Most recent pan-specific methods use deep learning (DL), such as Convolutional Neural Network (CNN), Long Short-Term Memory (LSTM) and transformer to learn the interaction between MHC-II molecules and peptides. There exist four representative DL-based methods: PUFFIN [13], DeepSeqPanII [14], MHCAttnNet [15] and BERTMHC [16]. In spite of using advanced DL techniques, these methods concatenate the sequence encoding of an MHC-II molecule and a peptide for the input, which do not take advantage of important domain knowledge, resulting in the lacking of performance improvement and interpretability.

We propose a novel deep learning based method, DeepMHCII, for accurate MHC-II pepitde binding affinity prediction by incorporating biological knowledge into designing the model architecture. Specifically DeepMHCII is modeled, considering the following three distinct features: 1) binding core and PFRs in each peptide; 2) pseudo sequence, i.e. the sequence with only crucial residues for directly interacting with the binding core of the counterpart peptide, in each MHC-II molecule. 3) interaction between a peptide and an MHC-II molecule by the interaction between all potential binding cores and the pseudo sequence. Note that these three features have not been explicitly addressed by any existing methods simultaneously. Specifically, DeepMHCII generates a binding interaction convolutional layer (BICL) with adaptive kernel size filters to model the interaction between peptides and MHC-II molecules.

The performance of DeepMHCII has been thoroughly validated by extensive experiments on four benchmark datasets under various settings, such as five fold cross validation, leave one molecule out (LOMO), independent testing set verification, and binding core prediction. We compared the predictive performance of DeepMHCII with four state-of-the-art methods: NetMHCIIpan-3.2 [6], PUFFIN [13], DeepSeqPanII [14] and MHCAttnNet [15]. Experimental results demonstrate that DeepMHCII outperformed all competing methods in all experiments. The improvement was especially significant in LOMO and independent testing set verification. For example, DeepMHCII achieved an average AUC of 0.77 in independent testing, which was 7% and 10% higher than that of NetMHCIIpan-3.2 (0.719) and PUFFIN (0.70). In addition, DeepMHCII achieved an average PCC (Pearson Correlation Coefficient) of 0.560 in LOMO, which was 6.7% higher than that of PUFFIN (0.525). All these results indicate the effectiveness of DeepMHCII on predicting the binding specificity of unseen MHC-II molecules. We also verified the performance advantage and interpretability of DeepMHCII in identifying the binding core of peptides and binding motifs of MHC-II molecules.

## 2 Method

### 2.1 Problem Formulation

Suppose *P* = (*p*_1_, *p*_2_, …, *p*_*L*_) denotes the peptide sequence and *Q* = (*q*_1_, *q*_2_, …, *q*_*L*′_) denotes the *L*′-length MHC-II molecule sequence, where each of *p*_*i*_ and *q*_*j*_ stands for one of 20 types of amino acids. The task is a regression problem to predict the binding affinity 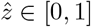 by a given pair of *P* and *Q*. The binding affinity is mainly determined by the nine-length binding core (unknown) of the peptide and the binding groove with nine pockets in the MHC-II molecule. In addition, it has been found that peptides flanking regions (PFRs) of binding core affect the binding affinity.

### 2.2 Overview

The basic idea of DeepMHCII is 1) to use deep learning to explicitly model the interaction between an MHC-II molecule sequence and a peptide, and 2) considering the crucial residue for binding, to focus on only the interaction between the binding core in a peptide and counterpart important amino acid residues (pseudo sequence) in a MHC-II molecule. Specifically, the input of DeepMHCII is a *L*-length peptide sequence *P* and, a 34-length pseudo sequence *Q*′ extracted from an MHC-II molecule sequence *Q* [17, 6] (More details about pseudo sequence are described in the **Appendix**). Figure 1 shows the architecture of DeepMHCII with mainly four steps: 1) we apply an embedding layer to the peptide sequence and also another embedding layer to the pseudo sequence of MHC-II to obtain deep semantic dense representations; 2) we use a binding interaction convolutional layer (BICL) to obtain the representation of binding interaction between the potential binding cores and the MHC-II molecule; 3) we use fully connected layers and a max-pooling layer to extract the interaction of peptide and MHC-II molecule; 4) we use an output layer to obtain the predicted binding affinity.

**Figure 1:**
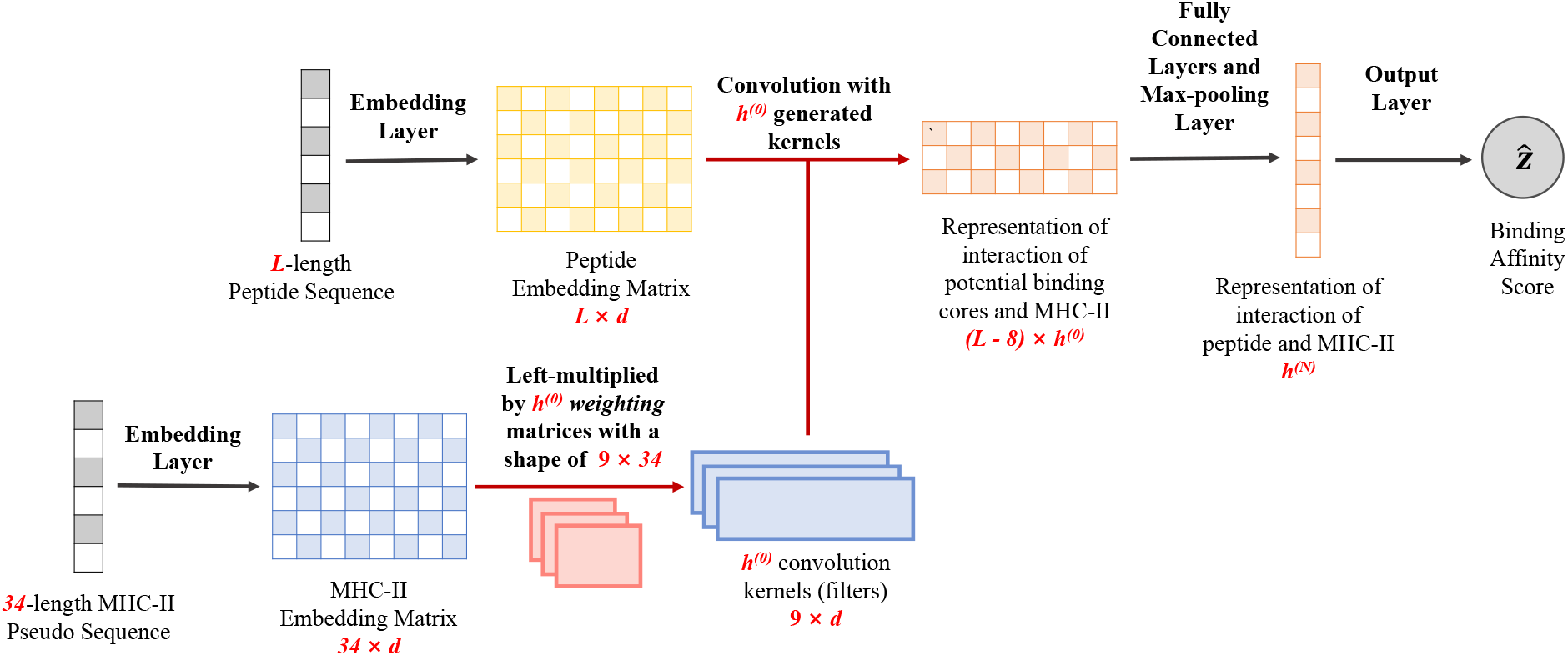
The architecture of DeepMHCII. The red arrows are processes of the binding interaction convolutional layer (BICL).

### 2.3 Input Layer

We use an embedding layer to encode amino acid sequences for peptides and also similarly another embedding layer for MHC-II pseudo sequences. Let *L* be the length of an input peptide and *d* be the dimension of amino acid embeddings. Then, for a given pair of a peptide sequence *P* and an MHC-II molecule pseudo sequence *Q*′, **X** ∈ ℝ^*L*×*d*^, the output of the embedding layer for *P*, and **Y** ∈ ℝ^34×*d*^, the output of the embedding layer for *Q*′, are given as follows:

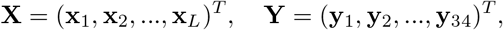

where **x**_*i*_ ∈ ℝ^*d*^ and **y**_*j*_ ∈ ℝ^*d*^ are the representation of the *i*-th amino acid of the peptide sequence and the *j*-th amino acid of the MHC-II pseudo sequence, respectively.

### 2.4 Binding Interaction Convolutional Layer

We use a binding interaction convolutional layer (BICL) to obtain the representation of the interaction between peptide **X** and MHC-II molecule **Y** by considering all possible binding cores of **X**. Traditionally in sequence-based CNN, input sequences share the same kernels (filters), while in our problem, each MHC-II molecule has a distinguished binding preference. To address this issue, BICL generates different kernels for each MHC-II molecule. Specifically, letting *M* be the size of a binding core(=9 in our problem), we use a weight matrix **W**_*k*_ with the size of *M* × 34 to generate the *k*-th kernel as *f* (**W**_*k*_**Y**), where *f* is the activation function. We can write a potential binding core of **X** starting with the *i*-th residue as **X**_*i*:*i*+*M*−1_. By using **X**_*i*:*i*+*M*−1_ and kernel *f* (**W**_*k*_**Y**), the interaction between potential binding core **X**_*i*:*i*+*M*−1_ and the MHC-II molecule **Y** can be given as follows:

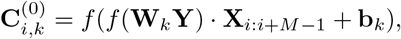

where **b**_*k*_ is the bias. Then the output of BICL can be written as 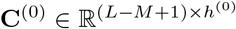, where *h*^(0)^ is the number of kernels.

Considering the effect of both binding core and PFRs, we used four different kernel sizes: 9, 11, 13 and 15. For example, kernel size of 15 is used to consider the effect of three more amino acids in the left and right side of binding core, respectively. For each kernel size, we used a different number of kernels. That is, the number of kernels for the kernel size of nine was the largest, followed by those of 11, 13 and 15. This setting will be described in detail in Section 3.2. Note that the padding symbol will be added to both side of **X** for kernel sizes other than 9 to get the same number of rows (number of potential binding cores) of output.

### 2.5 Max-pooling and Output Layer

We then use *N* fully connected layers and a max-pooling layer to obtain the representation 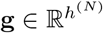 of the interaction between peptide **X** and MHC-II molecule **Y** as follows:

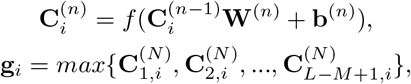

where 1 ≤ *n* ≤ *N*, and 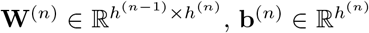 and 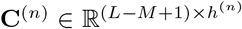 are the weights, bias and output of the *n*-th fully connected layer, respectively.

Finally, we use the output layer to obtain the predicted binding affinity 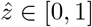 as follows:

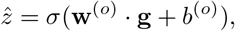

where 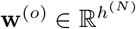 and *b*^(*o*)^ ∈ ℝ are the weights and bias, respectively. Our training objective is to minimize the mean square error (MSE). Practically we trained *T* models with different random initial weights and used the average over the *T* predicted scores as the final prediction.

### 2.6 Binding Core Prediction

For a given pair of peptide sequence and MHC-II molecule, we used each 9-length potential binding core and their PFR as the input of DeepMHCII, where the one with the highest score will be recommended as the binding core of this pair.

## 3 Experiments

### 3.1 Datasets

We used four publicly available benchmark datasets (BD2016, ID2017, BD2020 and BC2015) to train and evaluate DeepMHCII and other competing methods: BD2016 and ID2017 for MHC-peptide binding affinities, BD2020 for MHC-peptide binding classification and BC2015 for predicting binding cores.

BD2016^1^: BD2016 contains 134,281 data points of MHC-peptide binding affinities over 80 different MHC-II molecules, including 36 HLA-DR, 27 HLA-DQ, 9 HLA-DP and 8 H-2 molecules. BD2016 was compiled for training NetMHCIIpan-3.2 [6] from IEDB. The original, experimentally obtained IC50 binding value of each data point was transformed into the binding affinity with the range of [0,1] by 1 −log(IC_50_*nM*)*/* log(50, 000). BD2016 already provides five-fold cross-validation (5-fold CV) split which groups the peptides with common motifs into the same fold. Table 1 shows a summary of BD2016.

**Table 1:** Summary statistics of BD2016.

| Allele | #peptides | #binders | #MHCs |
| --- | --- | --- | --- |
| HLA-DR | 87,363 | 40,756 | 36 |
| HLA-DP | 15,564 | 5,135 | 9 |
| HLA-DQ | 28,081 | 9,098 | 27 |
| H-2 | 3,273 | 894 | 8 |
| Total | 134,281 | 55,883 | 80 |

ID2017: a dataset of MHC-peptide binding affinities was compiled from IEDB in 2017 for evaluating different MHC-II binding peptide prediction methods [18]. From this dataset, we generated an independent test dataset, ID2017, by removing data points overlapped with BD2016 and retaining MHC-II molecules with more than 50 peptides for robust performance evaluation. There are 10 HLA-DB molecules with 857 peptides in ID2017.

BD2020: a binary classification dataset of MHC-II peptide binding, which was extracted by Venkatesh et al. from IEDB for training MHCAttnNet [15]. Note that BD2020 has no quantitative binding affinities. BD2020 consists of 65,954 data points for 49 HLA-DRB molecules, where 36,035 are positive, 28,919 are negative and 5-fold CV split is also provided.

BC2015: a binding core benchmark, which was used to evaluate the performance of NetMHCIIpan-3.2 in identifying the binding core of an MHC-peptide complex. BC2015 consists of 51 complexes from PDB.

### 3.2 Experimental Settings

DeepMHCII used the following hyperparameter values, which were selected by 5-fold CV over BD2016: *d* (dimension of embeddings of amino acids) = 16. The numbers of kernels of BICL with the kernel sizes of 9, 11, 13 and 15 were 256, 128, 64 and 64, respectively. *N* (number of fully connected layers) = 2 and the sizes of nodes at the two layers were 256 and 128. *f* (activation function) was ReLU. We used batch normalization [19] after BICL and each of fully connected layers. Also we used dropout [20] with the drop rate of 0.25 to avoid overfitting. During the training process, the batch size was 128, the number of epochs was 20 and the optimizer we used was Adadelta [21] with the learning rate of 0.9 and weight decay of 1e-4. *T* (number of trained models) was 20.

We compared DeepMHCII with four state-of-the-art methods: NetMHCIIpan-3.2 ^2^ [6], PUFFIN^3^ [13], DeepSeqPanII^4^ [14] and MHCAttnNet^5^ [15]. All are neural network based methods. Since the training code of NetMHCIIpan-3.2 is unavailable, we used the experimental results (on BD2016) and the trained models provided by the authors directly. We trained PUFFIN and DeepSeqPanII on BD2016 using the implementation by the authors. According to the original paper and for a fair comparison, NetMHCIIpan-3.2 and both PUFFIN and DeepSeqPanII used a bagging ensemble of 200 and 20 models, respectively.

MHCAttnNet used the cross entropy objective function and binary MHC-peptide binding dataset (BD2020) for model training. We then trained DeepMHCII with the same dataset and the cross entropy objective function and compared the performance of MHCAttnNet obtained from the paper directly. Both DeepMHCII and MHCAttnNet used only one single model without any ensemble.

### 3.3 Evaluation Metrics

We set up binary classification: we used the area under the receiver operating characteristics curve (AUC) for each MHC-II molecule and reported the average AUC. Also to classify peptides into binders and non-binders, a binding threshold of 500nM was used: All peptides with an IC_50_ binding value < 500nM (0.426 after transformation) were classified as binders. Since there are a large number of MHC molecules, we used binomial test to check the statistical significance of performance difference (significance level was *p*-value < 0.05). In addition, we used the Pearson correlation coefficient (PCC) to examine the linear relationship between the predicted binding affinity and the true value.

### 3.4 Experimental Results

We first examined the performance of DeepMHCII and other competing methods by 5-fold CV over BD2016. To estimate the performance of different methods over unseen MHC-II molecule, following NetMHCIIpan-3.2, we conducted Leave-one-molecule-out (LOMO) over BD2016 by using the above 5-fold CV set-up. Specifically, each time, data points of only one MHC-II molecule in test fold were used for testing, while data points of all other MHC-II molecules in training folds were used for training over 5-fold CV settings. Following the settings in NetMHCIIpan-3.2, we focused on, out of all 81 MHC-II molecules, 61 molecules with more than 40 data points and at least 3 binders for the robustness of performance evaluation. We then explored performance comparison on ID2017, i.e. the independent test set.

#### 3.4.1 Comparison of DeepMHCII and Competing Methods under Five-Fold Cross-Validation

Table 2 reports the average AUC and PCC over all MHC-II molecules of 5-fold CV over BD2016 by DeepMHCII, NetMHCIIpan-3.2, PUFFIN and DeepSeqPanII (the detailed results of each MHC-II molecule are shown in the **Appendix**). DeepMHCII outperformed all competing methods in both AUC and PCC. For example, DeepMHCII achieved the highest AUC of 0.856, which was followed by NetMHCIIpan-3.2 (0.847), PUFFIN (0.846) and DeepSeqPanII (0.759). Also DeepMHCII outperformed NetMHCIIpan-3.2, PUFFIN and DeepSeqPanII on 48, 51 and 61, respectively, out of all 61 MHC-II molecules, all being statistically significant (two-tailed binomial test, *p*-value = 7.67 × 10^−6^, 9.62 × 10^−8^, 8.67 × 10^−19^, respectively).

**Table 2:** Performance of DeepMHCII and competing methods.

| Method | 5-CV |  | LOMO |  | IEBD Test | Binding Core |
| --- | --- | --- | --- | --- | --- | --- |
|  | AUC | PCC | AUC | PCC | AUC | #correct / #total |
| NetMHCIIpan-3.2 | 0.847 | 0.679 | 0.775 | 0.544 | 0.719 | 45/51 |
| PUFFIN | 0.846 | 0.676 | 0.768 | 0.525 | 0.700 | - |
| DeepSeqPanII | 0.759 | 0.524 | 0.732 | 0.473 | 0.700 | 10/51 |
| DeepMHCII | **0.856** | **0.691** | **0.785** | **0.560** | **0.770** | **47/51** |

#### 3.4.2 Comparison of DeepMHCII and Competing Methods under LOMO

Also Table 2 reports the average AUC and PCC of LOMO over all MHC-II molecules by DeepMHCII and competing methods (the detailed results of each MHC-II molecule are shown in the **Appendix**). The results are consistent with those of 5-fold CV. DeepMHCII achieved the highest AUC and PCC of 0.785 and 0.560, respectively, which were 1.3% and 2.9%, respectively, higher than those achieved by NetMHCIIpan-3.2, the second best method. Figure 2 plots the LOMO AUC results obtained by: DeepMHCII for y-axis and (a) NetMHCIIpan-3.2, (b) PUFFIN and (c) DeepSeqPanII for x-axis, where each dot corresponds to one MHC-II molecule. That is, if an MHC-II molecule appearing above the diagonal line, DeepMHCII outperformed the competing method. DeepMHCII outperformed both NetMCHIIpan-3.2, PUFFIN and DeepSeqPanII on 43, 42, 51 out of all 61 molecules, respectively, all being statistically significant (two-tailed binomial test, *p*-value = 1.87 × 10^−3^, 4.44 × 10^−3^ and 8.67 × 10^−19^, respectively). These results indicate that DeepMHCII is more robust and can deal with unknown MHC-II alleles better than the competing methods.

**Figure 2:**
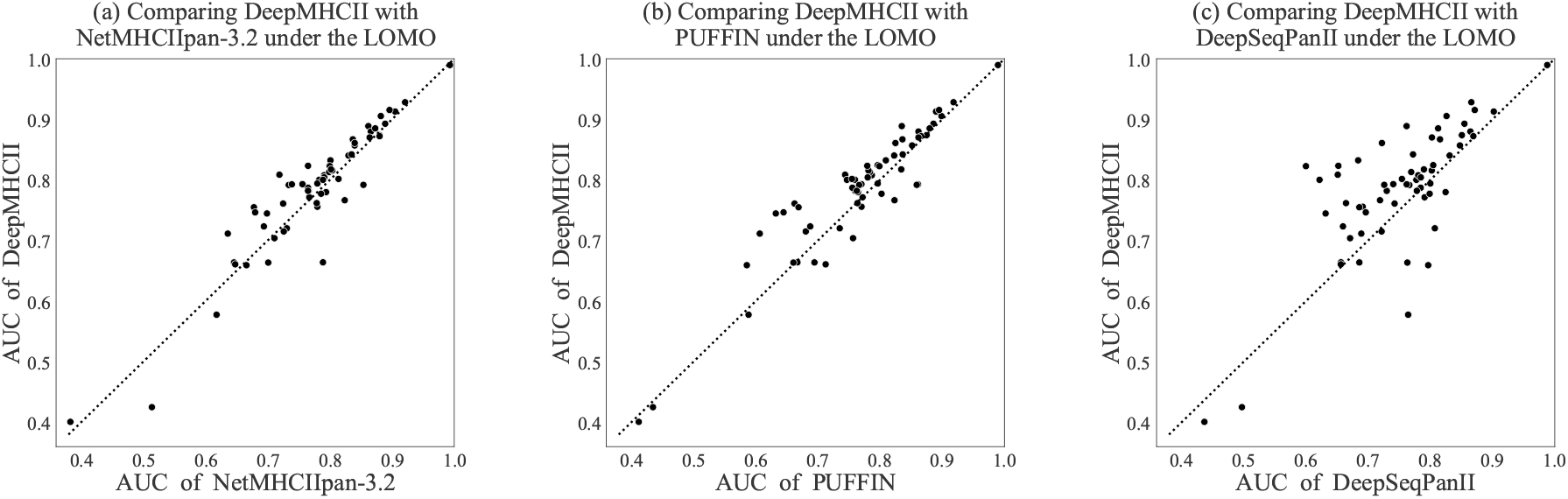
Performance comparison between DeepMHCII and (a) NetMHCIIpan-3.2, (b) PUFFIN and (c) DeepSeqPanII under LOMO. Each dot represents an MHC-II molecule.

#### 3.4.3 Comparison of DeepMHCII and Competing Methods on Independent Testing Set

Table 3 reports the performance on each MHC-II molecule of ID2017, the independent testing set, by DeepMHCII and competing methods. Figure 3 shows the ROC curves of DeepMHCII and competing methods on the whole ID2017. DeepMHCII outperformed all competing methods on both AUC of the whole ID2017 in Figure 3 and average AUC over all MHC-II molecules in Table 3. Specifically, DeepMHCII achieved the best average AUC of 0.770, which was 7.1% higher than the second best method, NetMHCIIpan-3.2 (0.719), and more than 10% higher than the other two competing methods, PUFFIN and DeepSeqPanII. Regarding the AUC of the whole testing set, DeepMHCII achieved the highest AUC of 0.775, which was was 7.8%, 12.0% and 15.5% higher than NetMHCIIpan-3.2 (0.719), PUFFIN (0.692) and DeepSeqPanII (0.671), respectively. For the performance over each MHC-II molecule, DeepMHCII outperformed NetMCHIIpan-3.2 and PUFFIN in all ten MHC-II molecules and DeepSeqPanII in eight out of the ten MHC-II molecules.

**Table 3:** Performance (AUC) of DeepMHCII and competing methods on the independent testing set.

| Allele | #peptides | #binders | NetMHCIIpan-3.2 | PUFFIN | DeepSeqPanII | DeepMHCII |
| --- | --- | --- | --- | --- | --- | --- |
| DBR1*01:01 | 100 | 81 | 0.880 | 0.834 | 0.733 | **0.882** |
| DBR1*03:01 | 99 | 61 | 0.588 | 0.620 | 0.536 | **0.629** |
| DBR1*04:01 | 142 | 91 | 0.787 | 0.762 | 0.748 | **0.863** |
| DBR1*07:01 | 94 | 72 | 0.800 | 0.728 | 0.732 | **0.814** |
| DBR1*09:01 | 62 | 75 | 0.793 | 0.796 | 0.654 | **0.889** |
| DBR1*11:01 | 94 | 72 | 0.614 | 0.629 | **0.699** | 0.657 |
| DBR1*12:02 | 59 | 48 | 0.661 | 0.742 | **0.824** | 0.788 |
| DBR1*13:01 | 57 | 47 | 0.538 | 0.472 | 0.591 | **0.615** |
| DBR1*15:01 | 96 | 81 | 0.767 | 0.683 | 0.751 | **0.799** |
| DBR1*15:02 | 54 | 41 | 0.760 | 0.735 | 0.730 | **0.764** |
| Average |  |  | 0.719 | 0.700 | 0.700 | **0.770** |

**Figure 3:**
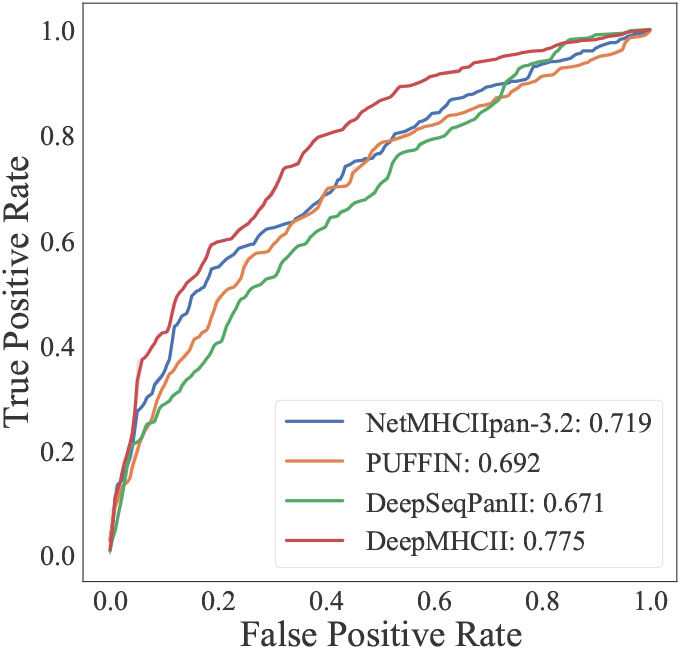
ROC curves by ID2017.

#### 3.4.4 Comparison with MHCAttnNet

We trained DeepMHCII with the cross entropy objective function under the same 5-fold CV split over BD2020 as MHCAttnNet, a binary MHC-peptide binding classification method. Table 4 shows the performance of DeepMHCII and MHCAttnNet, where following the original paper of MHCAttnNet, we used AUC over the whole test set, instead of the average AUC per MHC-II molecule. The AUC and accuracy of DeepMHCII were 4.1% and 3.0%, respectively, higher than MHCAttnNet, further demonstrating the performance advantage of DeepMHCII.

**Table 4:** Performance of DeepMHCII and MHCAttnNet on BD2020.

| Method | Accuracy | AUC |
| --- | --- | --- |
| MHCAttnNet | 0.755 | 0.758 |
| DeepMHCII | **0.786** | **0.781** |

### 3.5 Result Analysis

For examining the interpretability of DeepMHCII, we firstly compared DeepMHCII with DeepSeqPanII and NetMHCIIpan-3.2 in binding core prediction over BC2015. We then visualized the binding motifs of MHC-II molecules and compared the sequence logos generated by DeepMHCII, DeepSeqPanII, and NetMHCIIpan-3.2. Furthermore, to illustrate the biological perspective captured by DeepMHCII, we analyzed the weights of BICL in DeepMHCII. Finally, we explore the peformance of DeepMHCII with different settings of kernel size.

#### 3.5.1 Binding Core Prediction

Table 2 shows the results over BC2015 in the “Binding Core” column, where PUFFIN could not predict binding cores and thus could not be shown. The detailed results are shown in the Table A1 in **Appendix**. Note that NetMHCIIpan-3.2 has one variant, “NetMHCIIpan-3.2 (without offset)”, which uses the original prediction results to identify the binding core. Out of all 51 pairs, DeepMHCII correctly predicted 47, being followed by NetMHCIIpan-3.2 (45), NetMHCIIpan-3.2 without offest (28) and DeepSeqPanII (10). This result highlights the advantage of DeepMHCII of flexibly modeling the binding core and its high interpretability.

#### 3.5.2 Sequence logos

We visualized the binding motifs of MHC-II molecules obtained by each prediction method as sequence logos [22]. Following the description in [23], we first computed the binding scores of 100,000 random peptides from SwissProt and then selected the top 1% predicted binders to draw sequence logos (with default settings). Since PUFFIN does not have the ability to predict the binding core, we compared the sequence logos generated by DeepMHCII, NetMHCIIpan-3.2 and DeepSeqPanII. We focused on four MHC-II molecules, DRB1*04:01, DRB1*09:01, DRB1*12:02 and DRB1*13:01, in ID2017, where DeepMHCII outperformed NetMHCIIpan-3.2 most. Figure A1 in **Appendix** shows the sequence logos of these four MHC-II molecules by different methods. In each column, the total height represents the relative information content (also importance) of the corresponding position (pocket) in the motif, and the height of each letter (amino acid) shows the frequency of the corresponding amino acid in the position. It is widely observed and generally thought that P1 (pocket 1), P4, P6 and P9 are four primary anchors, which are most important for peptide binding [24]. All four sequence logos of DeepMHCII are consistent with this widely accepted understanding. In contrast, DeepSeqPanII could not distinguish these four primary anchors from other five pockets at all. Also the sequence logos by NetMHCIIpan3.2 contained noise at pockets, especially non primary pockets in DRB1*12:02 and DRB1*13:01. By taking a closer look, we could observe clear differences among prediction methods in amino acid preference in primary anchors. For example, P4 of DRB1*04:01 was identified as a primary anchor by both DeepMHCII and NetMHCIIpan-3.2, where preferred amino acids in P4 by DeepMHCII were [DILAES] but [LIASVDM] by NetMHCIIpan-3.2. According to SYFPEITHI [24], an MHC binding motif database, P4 allows amino acid E, being consistent with the sequence logo of DeepMHCII. Overall, DeepMHCII could generate a better sequence logo than the competing methods.

#### 3.5.3 Analyzing weights of BICL

We examined the importance of each position in the binding core by checking the absolute value of the weight (obtained by BICL) for each pocket of the binding core. For each pocket, we summed the absolute weight values for each model and computed the mean and standard deviation from all *T* = 20 models of DeepMHCII. Table 5 shows the mean and standard deviation of each pocket. The values of positions 1, 4, 6 and 9 were much larger than other positions, being consistent with that P1 (pocket 1), P4, P6 and P9 are primary anchors [24]. From this result, DeepMHCII learns biological knowledge from data, showing the validity of DeepMHCII from a biological perspective.

**Table 5:** Weights of BICL for nine pockets of the binding core.

| Pocket | Weight |
| --- | --- |
| P1 | **243.21**±10.14 |
| P2 | 230.51±10.22 |
| P3 | 232.19±8.48 |
| P4 | **256.36**±10.48 |
| P5 | 232.00±10.86 |
| P6 | **251.38**±10.02 |
| P7 | 239.53±9.44 |
| P8 | 227.72±9.80 |
| P9 | **253.76**±10.55 |

#### 3.5.4 Ablation Experiments on Kernels

We examined DeepMHCII with the single kernel size *k*, selecting *k* from 5, 7, 9, 11, 13 and 15, where we call DeepMHCII trained by the kernel size of *k* as DeepMHCII_*k*_ (Note that original DeepMHCII uses four different kernel sizes: 9, 11, 13 and 15 at once). Table 6 reports the performance of 5-fold CV over BD2016 by DeepMHCII_*k*_. We have three findings: 1) DeepMHCII_9_ outperformed DeepMHCII_5_ and DeepMHCII_7_ significantly. Specifically, DeepMHCII_9_ achieved PCC of 0.676, which was followed by DeepMHCII_7_ (0.651) and DeepMHCII_5_ (0.637). This is consistent with that the standard binding core is with nine amino acids. 2) DeepMHCII_11_, DeepMHCII_13_ and DeepMHCII_15_ achieved a similar performance, which was higher than DeepMHCII_9_. For example, both DeepMHCII_13_ and DeepMHCII_15_ achieved PCC of 0.686, which was higher than DeepMHCII_9_ (0.676). This suggests that PFR has some positive effect on MHC-II peptide binding. 3) DeepMHCII achieved the best performance with AUC of 0.856 and PCC of 0.691 among all compared methods. All these results confirm that the high performance of DeepMHCII was obtained by incorporating biological knowledge into the model design.

**Table 6:** Performance of DeepMHCII with different kernel sizes under 5-fold CV.

| Method | AUC | PCC |
| --- | --- | --- |
| DeepMHCII<sub>5</sub> | 0.828 | 0.637 |
| DeepMHCII<sub>7</sub> | 0.834 | 0.651 |
| DeepMHCII<sub>9</sub> | 0.849 | 0.676 |
| DeepMHCII<sub>11</sub> | 0.853 | 0.685 |
| DeepMHCII<sub>13</sub> | 0.854 | 0.686 |
| DeepMHCII<sub>15</sub> | 0.853 | 0.686 |
| DeepMHCII | **0.856** | **0.691** |

## 4 Conclusion

We have proposed a new deep learning model, DeepMHCII, for predicting peptide-MHC binding affinity, the binding core and important pockets in the binding core. Considering the biological characteristics of peptide-MHC binding, DeepMHCII have explicitly incorporated interaction processes between a peptide and an MHC-II molecule through interaction convolution layers, to biologically understand the peptide-MHC binding core and predict the binding affinity. Extensive experiments with four large-scale data sets demonstrate that DeepMHCII significantly outperformed all four state-of-the-art methods under various settings. Furthermore DeepMHCII captured the motifs more precisely than the compared methods, verifying the high performance in predicting the binding core and also the important pockets. All these results proved the usefulness of DeepMHCII in terms of the high accuracy but also precise biological discovery and high scientific interpretability. A limitation of our study is that we focus on peptide binding prediction other than epitope prediction, where the MHC binding peptide must be recognized by T cell receptor (TCR). Possible future work would be to incorporate more biological knowledge into model design to develop high-performance deep learning methods for epitope prediction [2].

## A Experimental Results

### A.1 Binding Core Prediction over BC2015

Table A1 shows the detailed results of DeepMHCII, DeepSeqPanII and NetMHCIIpan 3.2 over BC2015, where PUFFIN could not predict the binding core and it’s not be shown. For each pair (row) of an allele (MHC-II molecule) and antigen (peptide), red letters show the true binding core. Only the wrongly predicted binding core is shown, and the correctly predicted entry is blank, meaning that a method with a larger number of blank entries is better. Out of all 51 pairs, DeepMHCII correctly predicted 47, being followed by NetMHCIIpan-3.2 (45) and DeepSeqPanII (10).

**Table A1:**
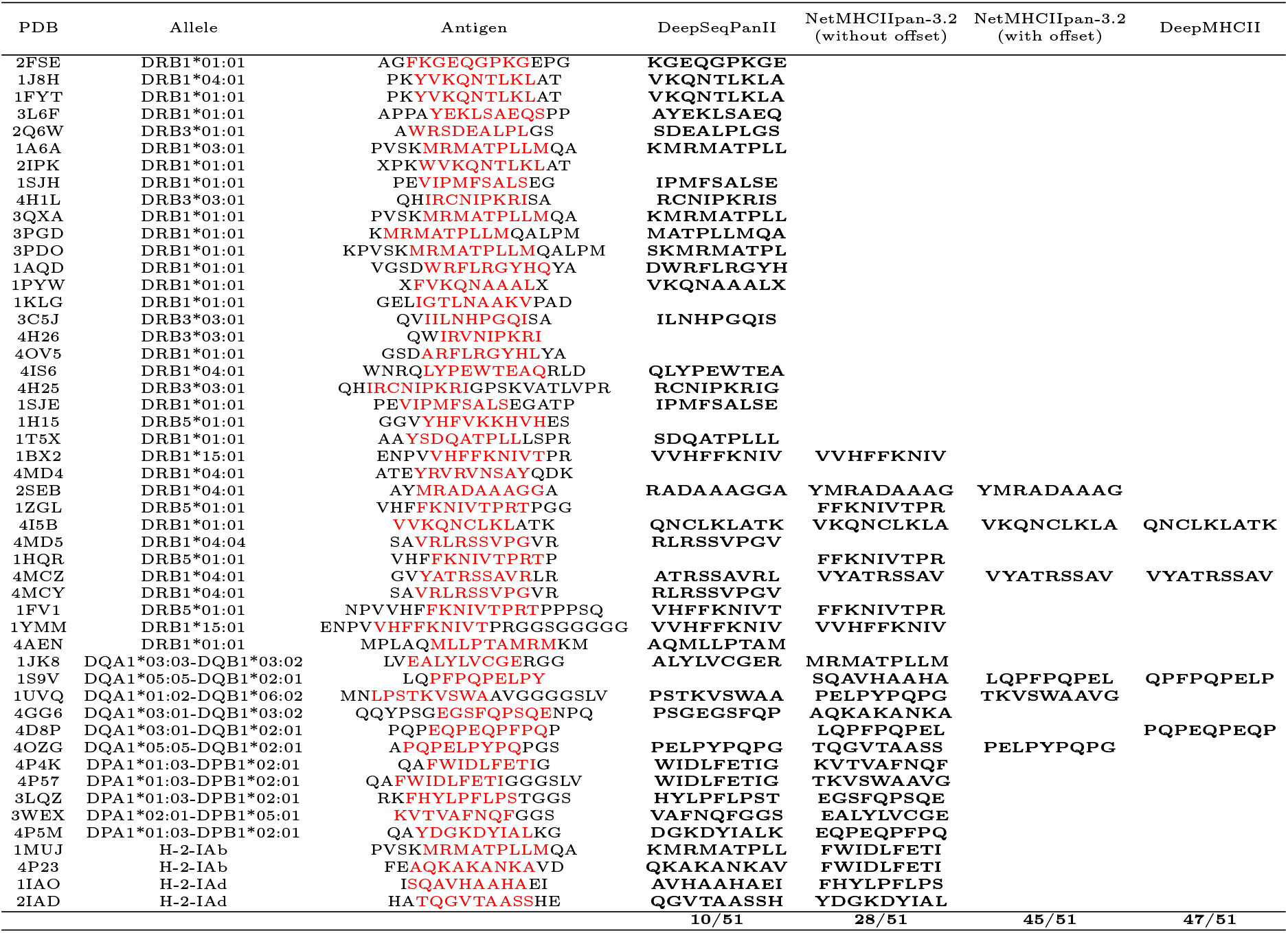
Binding core prediction results. Red letters are true cores. Only wrongly predicted cores are shown for each method.

### A.2 Sequence Logos over ID2017

We focused on four MHC-II molecules, DRB1*04:01, DRB1*09:01, DRB1*12:02 and DRB1*13:01, in ID2017, where DeepMHCII outperformed NetMHCIIpan-3.2 most. Figure A1 shows the sequence logos of these four MHC-II molecules by different methods. In each column, the total height represents the relative information content (also importance) of the corresponding position (pocket) in the motif, and the height of each letter (amino acid) shows the frequency of the corresponding amino acid in the position.

### A.3 Detailed Results of Five Fold Cross-Validation over BD2016

Table A2 shows the detailed results of DeepMHCII and competing methods under 5-CV for all MHC-II molecules. DeepMHCII achieved the best AUC and PCC on 45 and 48 out of 61 MHC-II molecules, respectively. In addition, DeepMHCII achieved the highest average AUC (0.856) and PCC (0.691).

**Figure A1:**
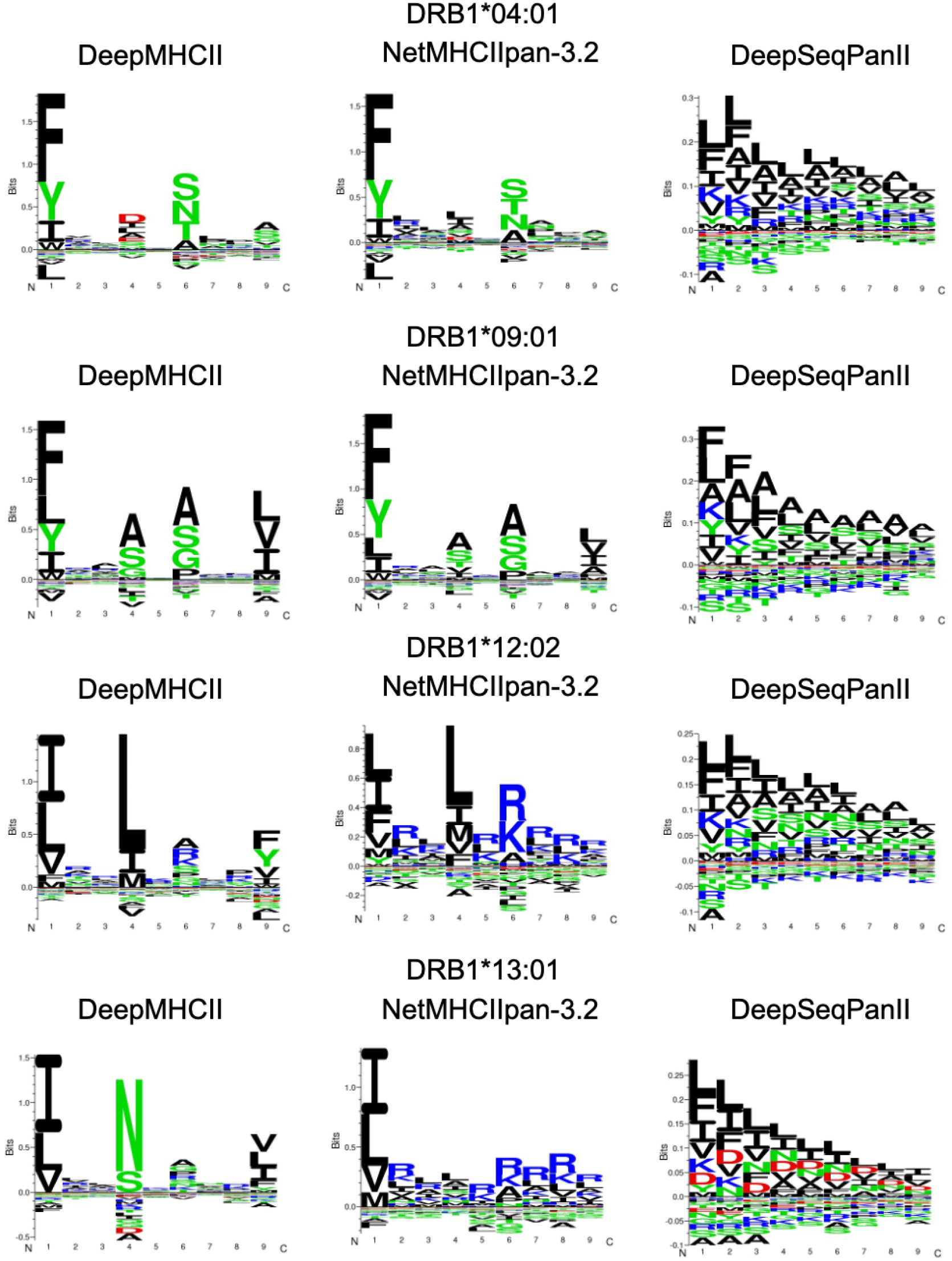
Sequence logos by DeepMHCII, NetMHCIIpan-3.2 and DeepSeqPanII.

### A.4 Detailed Results of LOMO over BD2016

Table A3 shows the detailed results of DeepMHCII and competing methods under LOMO for all MHC-II molecules. Similar to the results of five fold cross-validation, DeepMHCII achieved the the highest average AUC (0.817) and PCC (0.621). Specifically, DeepMHCII achieved the best AUC and PCC on 45 and 48 out of 61 MHC-II molecules, respectively.

**Table A2:** Detailed five-fold cross-validation performance of DeepMHCII and competing methods.

| Allele | NetMHCIIpan-3.2 |  | PUFFIN |  | DeepSeqPanII |  | DeepMHCII |  |
| --- | --- | --- | --- | --- | --- | --- | --- | --- |
|  | AUC | PCC | AUC | PCC | AUC | PCC | AUC | PCC |
| DRB1_0101 | 0.832 | 0.674 | 0.827 | 0.661 | 0.791 | 0.568 | **0.840** | **0.687** |
| DRB1_0103 | **0.678** | **0.408** | 0.658 | 0.364 | 0.566 | 0.387 | 0.638 | 0.217 |
| DRB1_0301 | 0.816 | 0.621 | 0.815 | 0.612 | 0.670 | 0.356 | **0.832** | **0.638** |
| DRB1_0401 | 0.809 | 0.613 | 0.805 | 0.600 | 0.730 | 0.467 | **0.818** | **0.629** |
| DRB1_0402 | **0.701** | **0.526** | 0.666 | 0.410 | 0.659 | 0.470 | 0.673 | 0.511 |
| DRB1_0403 | 0.841 | 0.615 | 0.805 | 0.568 | 0.737 | 0.461 | **0.897** | **0.706** |
| DRB1_0404 | 0.812 | 0.652 | 0.811 | 0.643 | 0.768 | 0.568 | **0.818** | **0.663** |
| DRB1_0405 | 0.827 | 0.650 | 0.832 | 0.660 | 0.757 | 0.518 | **0.847** | **0.685** |
| DRB1_0701 | 0.875 | 0.733 | 0.869 | 0.723 | 0.827 | 0.634 | **0.887** | **0.759** |
| DRB1_0801 | 0.844 | 0.671 | 0.856 | 0.687 | 0.800 | 0.588 | **0.860** | **0.693** |
| DRB1_0802 | 0.834 | 0.663 | 0.833 | 0.658 | 0.803 | 0.594 | **0.843** | **0.676** |
| DRB1_0901 | 0.833 | 0.666 | 0.832 | 0.658 | 0.787 | 0.571 | **0.841** | **0.678** |
| DRB1_1001 | 0.923 | 0.761 | 0.918 | 0.770 | 0.900 | 0.680 | **0.923** | **0.781** |
| DRB1_1101 | 0.864 | 0.728 | 0.859 | 0.727 | 0.799 | 0.594 | **0.873** | **0.746** |
| DRB1_1201 | 0.868 | 0.727 | 0.880 | 0.752 | 0.805 | 0.622 | **0.890** | **0.768** |
| DRB1_1301 | **0.857** | **0.658** | 0.843 | 0.649 | 0.794 | 0.528 | 0.821 | 0.612 |
| DRB1_1302 | 0.885 | 0.761 | 0.884 | 0.757 | 0.757 | 0.555 | **0.885** | **0.764** |
| DRB1_1501 | 0.834 | 0.666 | 0.838 | 0.672 | 0.789 | 0.571 | **0.847** | **0.693** |
| DRB1_1602 | 0.883 | 0.767 | 0.894 | **0.788** | 0.857 | 0.672 | **0.895** | 0.787 |
| DRB3_0101 | 0.888 | 0.787 | **0.904** | 0.803 | 0.751 | 0.521 | 0.903 | **0.804** |
| DRB3_0202 | 0.869 | 0.793 | 0.880 | 0.800 | 0.742 | 0.558 | **0.888** | **0.812** |
| DRB3_0301 | 0.840 | 0.665 | 0.833 | 0.649 | 0.765 | 0.506 | **0.845** | **0.705** |
| DRB4_0101 | 0.822 | 0.638 | 0.826 | 0.651 | 0.731 | 0.493 | **0.841** | **0.682** |
| DRB4_0103 | **0.841** | **0.629** | 0.822 | 0.588 | 0.806 | 0.562 | 0.839 | 0.612 |
| DRB5_0101 | 0.849 | 0.709 | 0.852 | 0.717 | 0.764 | 0.548 | **0.864** | **0.735** |
| H-2-IAb | 0.894 | 0.720 | 0.882 | 0.706 | 0.753 | 0.470 | **0.898** | **0.728** |
| H-2-IAd | 0.819 | 0.658 | 0.796 | 0.623 | 0.752 | 0.539 | **0.839** | **0.678** |
| H-2-IaK | **0.635** | 0.222 | 0.498 | **0.240** | 0.435 | -0.030 | 0.541 | 0.218 |
| H-2-IAs | **0.825** | **0.587** | 0.772 | 0.496 | 0.479 | 0.112 | 0.814 | 0.580 |
| H-2-IAu | 0.765 | 0.545 | 0.822 | **0.574** | 0.786 | 0.430 | **0.822** | 0.570 |
| H-2-IEd | 0.754 | **0.560** | **0.774** | 0.528 | 0.686 | 0.424 | 0.740 | 0.510 |
| H-2-IEk | **0.853** | **0.768** | 0.846 | 0.759 | 0.741 | 0.582 | 0.821 | 0.736 |
| HLA-DPA10103-DPB10201 | 0.917 | 0.741 | **0.921** | 0.736 | 0.832 | 0.583 | 0.919 | **0.755** |
| HLA-DPA10103-DPB10301 | 0.902 | 0.783 | 0.894 | 0.771 | 0.842 | 0.677 | **0.910** | **0.792** |
| HLA-DPA10103-DPB10401 | 0.935 | 0.884 | 0.933 | 0.881 | 0.879 | 0.775 | **0.942** | **0.892** |
| HLA-DPA10103-DPB10402 | 0.710 | 0.425 | 0.738 | 0.395 | 0.639 | 0.415 | **0.809** | **0.519** |
| HLA-DPA10103-DPB10601 | 0.995 | **0.962** | 0.993 | 0.955 | 0.972 | 0.912 | **0.995** | 0.957 |
| HLA-DPA10201-DPB10101 | 0.903 | 0.844 | 0.900 | 0.843 | 0.854 | 0.754 | **0.912** | **0.854** |
| HLA-DPA10201-DPB10501 | 0.911 | 0.804 | 0.911 | 0.804 | 0.874 | 0.741 | **0.914** | **0.810** |
| HLA-DPA10201-DPB11401 | 0.930 | 0.836 | 0.937 | 0.851 | 0.881 | 0.744 | **0.946** | **0.860** |
| HLA-DPA10301-DPB10402 | 0.904 | 0.837 | 0.903 | 0.836 | 0.856 | 0.754 | **0.910** | **0.844** |
| HLA-DQA10101-DQB10501 | 0.900 | 0.781 | 0.909 | 0.798 | 0.724 | 0.467 | **0.911** | **0.799** |
| HLA-DQA10102-DQB10501 | 0.839 | 0.654 | **0.849** | **0.660** | 0.751 | 0.491 | 0.824 | 0.627 |
| HLA-DQA10102-DQB10502 | 0.835 | 0.654 | 0.857 | 0.665 | 0.720 | 0.460 | **0.859** | **0.685** |
| HLA-DQA10102-DQB10602 | 0.890 | 0.798 | 0.877 | 0.779 | 0.827 | 0.679 | **0.906** | **0.823** |
| HLA-DQA10103-DQB10603 | 0.861 | 0.577 | 0.851 | 0.578 | 0.803 | 0.463 | **0.883** | **0.614** |
| HLA-DQA10104-DQB10503 | 0.805 | 0.596 | 0.830 | 0.620 | 0.686 | 0.405 | **0.843** | **0.666** |
| HLA-DQA10201-DQB10202 | 0.814 | 0.614 | 0.837 | 0.625 | 0.642 | 0.408 | **0.850** | **0.663** |
| HLA-DQA10201-DQB10301 | 0.849 | 0.649 | 0.852 | 0.660 | 0.795 | 0.561 | **0.873** | **0.695** |
| HLA-DQA10201-DQB10303 | 0.894 | 0.736 | 0.891 | 0.724 | 0.815 | 0.593 | **0.900** | **0.755** |
| HLA-DQA10201-DQB10402 | 0.860 | 0.634 | 0.861 | 0.642 | 0.714 | 0.373 | **0.875** | **0.656** |
| HLA-DQA10301-DQB10301 | **0.839** | **0.664** | 0.814 | 0.623 | 0.735 | 0.501 | 0.815 | 0.636 |
| HLA-DQA10301-DQB10302 | 0.810 | 0.657 | 0.847 | 0.697 | 0.650 | 0.388 | **0.847** | **0.700** |
| HLA-DQA10303-DQB10402 | 0.820 | 0.543 | 0.837 | **0.601** | 0.671 | 0.262 | **0.859** | 0.592 |
| HLA-DQA10401-DQB10402 | 0.883 | 0.759 | **0.893** | **0.781** | 0.813 | 0.609 | 0.879 | 0.746 |
| HLA-DQA10501-DQB10201 | 0.876 | 0.759 | 0.881 | 0.770 | 0.780 | 0.585 | **0.883** | **0.778** |
| HLA-DQA10501-DQB10301 | 0.915 | 0.786 | 0.913 | 0.783 | 0.758 | 0.488 | **0.928** | **0.815** |
| HLA-DQA10501-DQB10302 | 0.822 | 0.638 | **0.849** | **0.665** | 0.792 | 0.542 | 0.844 | 0.650 |
| HLA-DQA10501-DQB10303 | 0.876 | 0.682 | 0.875 | 0.706 | 0.820 | 0.566 | **0.887** | **0.712** |
| HLA-DQA10501-DQB10402 | 0.868 | **0.692** | **0.871** | 0.691 | 0.713 | 0.380 | 0.870 | 0.684 |
| HLA-DQA10601-DQB10402 | **0.848** | 0.568 | 0.831 | **0.579** | 0.671 | 0.264 | 0.836 | 0.530 |
| Average | 0.847 | 0.679 | 0.846 | 0.676 | 0.759 | 0.524 | **0.856** | **0.691** |
| p-value | 7.67e-6 |  | 4.59e-7 |  | 8.67e-19 |  |  |  |

**Table A3:** Detailed LOMO performance of DeepMHCII and competing methods.

| Allele | NetMHCIIpan-3.2 |  | PUFFIN |  | DeepSeqPanII |  | DeepMHCII |  |
| --- | --- | --- | --- | --- | --- | --- | --- | --- |
|  | AUC | PCC | AUC | PCC | AUC | PCC | AUC | PCC |
| DRB1_0101 | 0.783 | 0.558 | 0.760 | 0.511 | 0.772 | 0.527 | **0.800** | **0.600** |
| DRB1_0103 | 0.711 | **0.458** | **0.757** | 0.379 | 0.553 | 0.422 | 0.704 | 0.260 |
| DRB1_0301 | 0.699 | 0.400 | 0.632 | 0.272 | 0.630 | 0.279 | **0.745** | **0.481** |
| DRB1_0401 | 0.766 | 0.517 | 0.760 | 0.509 | 0.714 | 0.435 | **0.782** | **0.558** |
| DRB1_0402 | **0.789** | **0.646** | 0.667 | 0.415 | 0.656 | 0.413 | 0.664 | 0.498 |
| DRB1_0403 | 0.862 | 0.679 | 0.835 | 0.660 | 0.749 | 0.520 | **0.889** | **0.725** |
| DRB1_0404 | 0.791 | 0.598 | 0.787 | 0.583 | 0.762 | 0.554 | **0.808** | **0.631** |
| DRB1_0405 | 0.799 | 0.596 | 0.784 | 0.576 | 0.751 | 0.506 | **0.813** | **0.620** |
| DRB1_0701 | 0.830 | 0.634 | 0.823 | 0.628 | 0.816 | 0.611 | **0.840** | **0.666** |
| DRB1_0801 | 0.804 | 0.600 | 0.781 | 0.541 | 0.791 | 0.571 | **0.816** | **0.603** |
| DRB1_0802 | 0.765 | 0.510 | 0.756 | 0.500 | **0.794** | **0.570** | 0.787 | 0.556 |
| DRB1_0901 | 0.791 | 0.586 | 0.780 | 0.561 | 0.774 | 0.543 | **0.804** | **0.612** |
| DRB1_1001 | 0.905 | 0.698 | 0.890 | 0.699 | 0.894 | 0.664 | **0.913** | **0.743** |
| DRB1_1101 | 0.767 | 0.507 | 0.772 | 0.527 | **0.782** | **0.561** | 0.771 | 0.531 |
| DRB1_1201 | 0.800 | 0.592 | 0.796 | 0.609 | 0.757 | 0.571 | **0.824** | **0.653** |
| DRB1_1301 | 0.731 | 0.413 | 0.735 | 0.431 | **0.780** | **0.494** | 0.720 | 0.400 |
| DRB1_1302 | 0.701 | 0.437 | 0.661 | 0.340 | **0.728** | **0.510** | 0.664 | 0.354 |
| DRB1_1501 | 0.780 | 0.551 | **0.796** | 0.579 | 0.784 | 0.558 | 0.794 | **0.585** |
| DRB1_1602 | 0.866 | 0.731 | 0.862 | 0.711 | 0.858 | 0.665 | **0.880** | **0.752** |
| DRB3_0101 | 0.801 | 0.583 | 0.809 | 0.639 | 0.641 | 0.299 | **0.832** | **0.666** |
| DRB3_0202 | 0.756 | 0.596 | 0.770 | 0.600 | 0.738 | 0.528 | **0.793** | **0.649** |
| DRB3_0301 | 0.734 | 0.431 | 0.766 | 0.516 | 0.756 | 0.479 | **0.792** | **0.589** |
| DRB4_0101 | **0.726** | **0.462** | 0.681 | 0.390 | 0.703 | 0.451 | 0.715 | 0.452 |
| DRB4_0103 | 0.794 | 0.513 | 0.764 | 0.460 | **0.802** | **0.548** | 0.780 | 0.483 |
| DRB5_0101 | 0.765 | 0.553 | 0.763 | 0.538 | 0.751 | 0.525 | **0.782** | **0.578** |
| H-2-IAb | **0.780** | **0.557** | 0.770 | 0.544 | 0.681 | 0.348 | 0.756 | 0.487 |
| H-2-IAd | 0.725 | 0.461 | 0.663 | 0.334 | 0.724 | 0.490 | **0.761** | **0.523** |
| H-2-IaK | 0.383 | -0.102 | 0.412 | -0.080 | **0.426** | **-0.074** | 0.401 | -0.092 |
| H-2-IAs | **0.514** | **0.195** | 0.435 | 0.066 | 0.476 | 0.085 | 0.425 | 0.007 |
| H-2-IAu | 0.739 | 0.475 | **0.861** | **0.544** | 0.774 | 0.410 | 0.793 | 0.534 |
| H-2-IEd | 0.646 | 0.356 | 0.695 | 0.401 | **0.699** | **0.407** | 0.664 | 0.385 |
| H-2-IEk | 0.854 | 0.716 | **0.859** | **0.716** | 0.733 | 0.555 | 0.792 | 0.660 |
| HLA-DPA10103-DPB10201 | 0.882 | 0.669 | 0.899 | 0.696 | 0.810 | 0.536 | **0.905** | **0.716** |
| HLA-DPA10103-DPB10301 | 0.840 | 0.672 | 0.852 | 0.700 | 0.790 | 0.600 | **0.857** | **0.697** |
| HLA-DPA10103-DPB10401 | 0.921 | 0.863 | 0.918 | 0.858 | 0.851 | 0.723 | **0.928** | **0.872** |
| HLA-DPA10103-DPB10402 | 0.719 | 0.462 | 0.744 | 0.432 | 0.627 | 0.396 | **0.809** | **0.519** |
| HLA-DPA10103-DPB10601 | **0.993** | 0.945 | 0.990 | 0.943 | 0.971 | 0.909 | 0.990 | **0.950** |
| HLA-DPA10201-DPB10101 | **0.880** | **0.815** | 0.875 | 0.815 | 0.833 | 0.710 | 0.874 | 0.807 |
| HLA-DPA10201-DPB10501 | **0.880** | **0.756** | 0.866 | 0.736 | 0.864 | 0.715 | 0.872 | 0.750 |
| HLA-DPA10201-DPB11401 | 0.896 | 0.751 | 0.895 | 0.764 | 0.821 | 0.645 | **0.915** | **0.775** |
| HLA-DPA10301-DPB10402 | 0.889 | 0.805 | 0.886 | 0.807 | 0.842 | 0.727 | **0.893** | **0.816** |
| HLA-DQA10101-DQB10501 | 0.678 | 0.396 | 0.669 | 0.360 | 0.606 | 0.239 | **0.755** | **0.525** |
| HLA-DQA10102-DQB10501 | 0.618 | 0.224 | 0.589 | 0.181 | **0.740** | **0.474** | 0.578 | 0.175 |
| HLA-DQA10102-DQB10502 | 0.694 | 0.398 | 0.688 | 0.399 | 0.652 | 0.349 | **0.723** | **0.466** |
| HLA-DQA10102-DQB10602 | 0.786 | 0.561 | **0.802** | **0.616** | 0.776 | 0.569 | 0.777 | 0.573 |
| HLA-DQA10103-DQB10603 | 0.837 | 0.508 | 0.836 | 0.526 | 0.786 | 0.443 | **0.867** | **0.559** |
| HLA-DQA10104-DQB10503 | 0.765 | 0.534 | 0.779 | 0.555 | 0.628 | 0.321 | **0.823** | **0.631** |
| HLA-DQA10201-DQB10202 | 0.789 | 0.567 | 0.747 | 0.506 | 0.589 | 0.336 | **0.800** | **0.605** |
| HLA-DQA10201-DQB10301 | 0.835 | 0.620 | 0.836 | 0.625 | 0.749 | 0.468 | **0.842** | **0.641** |
| HLA-DQA10201-DQB10303 | 0.873 | 0.714 | 0.880 | 0.704 | 0.808 | 0.569 | **0.885** | **0.733** |
| HLA-DQA10201-DQB10402 | 0.840 | 0.590 | 0.825 | 0.563 | 0.707 | 0.357 | **0.861** | **0.628** |
| HLA-DQA10301-DQB10301 | **0.814** | **0.628** | 0.755 | 0.508 | 0.725 | 0.482 | 0.802 | 0.594 |
| HLA-DQA10301-DQB10302 | 0.648 | 0.379 | **0.713** | **0.481** | 0.603 | 0.308 | 0.661 | 0.405 |
| HLA-DQA10303-DQB10402 | 0.680 | 0.268 | 0.645 | 0.198 | 0.675 | 0.273 | **0.747** | **0.393** |
| HLA-DQA10401-DQB10402 | 0.666 | 0.297 | 0.586 | 0.169 | **0.789** | **0.552** | 0.659 | 0.290 |
| HLA-DQA10501-DQB10201 | **0.779** | **0.591** | 0.764 | 0.572 | 0.662 | 0.363 | 0.762 | 0.574 |
| HLA-DQA10501-DQB10301 | 0.800 | 0.575 | 0.799 | 0.578 | 0.582 | 0.160 | **0.823** | **0.615** |
| HLA-DQA10501-DQB10302 | 0.802 | 0.610 | 0.834 | 0.626 | 0.779 | 0.506 | **0.817** | **0.631** |
| HLA-DQA10501-DQB10303 | 0.864 | 0.677 | 0.862 | 0.673 | 0.797 | 0.532 | **0.870** | **0.684** |
| HLA-DQA10501-DQB10402 | **0.824** | **0.609** | 0.823 | 0.596 | 0.692 | 0.341 | 0.767 | 0.507 |
| HLA-DQA10601-DQB10402 | 0.636 | 0.181 | 0.607 | 0.119 | 0.662 | 0.248 | **0.711** | **0.302** |
| Average | 0.775 | 0.544 | 0.768 | 0.525 | 0.732 | 0.473 | **0.785** | **0.560** |
| p-value | 1.87e-3 |  | 9.62e-8 |  | 4.44e-3 |  |  |  |

### A.5 Running Time

Table A4 shows the training time and prediction speed of DeepMHCII and competing methods. All methods run on a single Nvidia Titan X (pascal). The training time of DeepMHCII is close to that of PUFFIN and much less than that of DeepSeqPanII which is based on RNN. Furthermore, DeepMHCII is much faster than both PUFFIN and DeepSeqPanII in prediction. The model size of DeepMHCII is also much smaller than both PUFFIN and DeepSeqPanII.

**Table A4:** Computation time and model size of DeepMHCII and competing methods.

| Method | Training (s) | Test (ms/sample) | Model Size (MB) |
| --- | --- | --- | --- |
| PUFFIN | 651 | 2.252 | 24.1 |
| DeepSeqPanII | 7,967 | 0.828 | 15.3 |
| DeepMHCII | 693 | 0.583 | 1.4 |

## B Pseudo Sequence of MHC-II molecules

Pseudo Sequence of MHC-II refers to the important MHC residues that are considered being crucial for peptide binding. The 34-length MHC-II molecule pseudo sequence was first proposed by Karosiene et al. in NetMHCIIpan-3.0 [1], which was composed of 15 and 19 amino acid residues from *α* and *β* chains of MHC-II, respectively. These residues were extracted from MHC II-peptide complexes in PDB [2], which were polymorphic in MHC molecules and found in close contact (<4 *A*°) with peptide binding core in at least one MHC II-peptide complex. The 34-length MHC-II pseudo sequence has also been used in NetMHCIIpan-3.1 and 3.2 [3, 4]. Similarly, we use this pseudo sequence as the representation of MHC-II molecules.

## Footnotes

1 http://www.cbs.dtu.dk/suppl/immunology/NetMHCIIpan-3.2

2 http://www.cbs.dtu.dk/services/NetMHCIIpan-3.2

3 https://github.com/gifford-lab/PUFFIN

4 https://github.com/pcpLiu/DeepSeqPanII

5 https://github.com/gopuvenkat/MHCAttnNet

## Notes

### Competing Interest Statement

The authors have declared no competing interest.

